# Step-wise increases in *FT1* expression regulate seasonal progression of flowering in wheat (*Triticum aestivum* L.)

**DOI:** 10.1101/2020.06.30.181396

**Authors:** Adam Gauley, Scott A. Boden

## Abstract

Flowering is regulated by genes that respond to changing daylengths and temperature, which have been well-studied using controlled conditions; however, the molecular processes underpinning flowering in nature remain poorly understood. Here, we investigate the genetic pathways that coordinate flowering and inflorescence development of wheat as daylengths extend naturally in the field, using lines that contain variant alleles for the key photoperiod gene, *Photoperiod-1* (*Ppd-1*). We found flowering involves a step-wise increase in the expression of *FLOWERING LOCUS T1* (*FT1*), which initiates under day-neutral conditions of early spring. The incremental rise in *FT1* expression is overridden in plants that contain a photoperiod-insensitive allele of *Ppd-1*, which hastens the completion of spikelet development and accelerates flowering time. The accelerated inflorescence development of photoperiod-insensitive lines is promoted by advanced seasonal expression of floral meristem identity genes. The completion of spikelet formation is promoted by *FLOWERING LOCUS T2*, which regulates spikelet number and is activated by *Ppd-1*. In wheat, flowering under natural photoperiods is regulated by step-wise increases in the expression of *FT1*, which responds dynamically to extending daylengths to promote early inflorescence development. This research provides a strong foundation to improve yield potential by fine-tuning the photoperiod-dependent control of inflorescence development.

## INTRODUCTION

Plants use environmental signals including daylength and temperature to determine the optimal time to flower. By monitoring changes in daylength, plants flower with remarkable seasonal precision despite variable environmental conditions such as fluctuating daily temperatures. Temperate cereals such as bread wheat (*Triticum aestivum*) perceive the extending daylengths of spring to promote flowering so that seed production occurs under favourable conditions (Worland *et al.*, 1998; Fjellheim *et al.*, 2014). During domestication, alleles that modify the plant’s response to these seasonal cues have been used to expand the geographical range of cultivation and improve productivity in marginal environments – these alleles typically accelerate flowering by reducing the requirement for long days or cold temperatures (Danyluk *et al.*, 2003; Trevaskis *et al.*, 2003; Yan *et al.*, 2003; Yan *et al.*, 2004; Turner *et al.*, 2005; Beales *et al.*, 2007).

In wheat, the responsiveness to daylength (photoperiod) is largely determined by allelic diversity for *Photoperiod-1* (*Ppd-1*) (Laurie *et al.*, 1995; Turner *et al.*, 2005; Beales *et al.*, 2007). *Ppd-1* influences flowering by modifying the expression of *FLOWERING LOCUS T1* (*FT1*), which is a conserved activator of flowering in plants (Turner *et al.*, 2005; Yan *et al.*, 2006; Beales *et al.*, 2007; Wilhelm *et al.*, 2009; Kitagawa *et al.*, 2012; Boden *et al.*, 2015; Bratzel & Turck, 2015). FT1 protein is expressed in leaves and transported to the shoot apical meristem (SAM), where it forms a complex with FLOWERING LOCUS D-LIKE (FDL) and 14-3-3 proteins (Corbesier *et al.*, 2007; Tamaki *et al.*, 2007; Li & Dubcovsky, 2008; Taoka *et al.*, 2011). The complex activates expression of meristem identity genes, which promote reproductive development of the inflorescence meristem (IM) (Corbesier *et al.*, 2007; Tamaki *et al.*, 2007; Li & Dubcovsky, 2008). In hexaploid wheat, photoperiod-insensitive alleles of *Ppd-1* activate *FT1* expression in the absence of long-day photoperiods – these alleles carry deletions in the *cis*-regulatory regions of *Ppd-1* or additional copies of the gene on the A, B and D genomes (termed *Ppd-A1a, Ppd-B1a* and *Ppd-D1a*, respectively) (Beales *et al.*, 2007; Diaz *et al.*, 2012; Kitagawa *et al.*, 2012; Shaw *et al.*, 2012; Seki *et al.*, 2013; Shaw *et al.*, 2013). Under constant short daylengths, the *cis*-regulatory mutations alter the daily rhythms of *Ppd-1* expression, causing it to be expressed in the evening when it is otherwise suppressed. While photoperiod-insensitive alleles confer early flowering phenotypes under all growth conditions, our understanding about the molecular function of *Ppd-1* comes from experiments performed in controlled growth environments (Beales *et al.*, 2007; Wilhelm *et al.*, 2009; Diaz *et al.*, 2012; Kitagawa *et al.*, 2012; Shaw *et al.*, 2012; Shaw *et al.*, 2013; Boden *et al.*, 2015). These experiments used extreme short (8-9 hours) or long (15-16 h) daylengths, which are different from the photoperiods when plants initiate flowering naturally in the field (11-13 h). These analyses have also focused on the role of *Ppd-1* in leaves, with little attention given to the impact of photoperiod-insensitive alleles on expression of meristem identity genes in the developing inflorescence. Given photoperiod-insensitive alleles reduce yield potential by significantly decreasing spikelet number and the survival of floret primordia (i.e. floret fertility), it is vital that we learn more about the role of *Ppd-1* in inflorescences of field-grown plants under natural daylengths to devise strategies for improving grain production (González *et al.*, 2005; Fischer *et al.*, 2014; González-Navarro *et al.*, 2015; Prieto *et al.*, 2018; Perez-Gianmarco *et al.*, 2019).

To understand how flowering is regulated in a seasonal context, we investigated the photoperiod-dependent flowering pathway and early inflorescence development under natural photoperiods in the field. We used near-isogenic lines (NILs) containing photoperiod-insensitive, sensitive and null alleles of *Ppd-1* to genetically alter the ability of wheat to perceive changes in daylength – the insensitive *Ppd-D1a* allele represents the majority of wheat grown in spring-type mega-environments (Shaw *et al.*, 2013). Our work demonstrates that the floral-promoting pathway of wheat dynamically responds to increasing daylengths to regulate spikelet and floret development, and presents new genetic targets for yield improvement.

## MATERIALS AND METHODS

### Plant material and growth conditions

Hexaploid wheat (*Triticum aestivum*) used here included: wild-type photoperiod-sensitive *cv.* Paragon; Paragon NILs containing the *Ppd-D1a* photoperiod-insensitive allele (Shaw *et al.*, 2013) or null *ppd-1* alleles on the A, B and D genomes (Shaw *et al.*, 2013); two null *ft-b2* mutants (*Cad0122* and *Cad1655*) obtained from the hexaploid wheat TILLING population (**S9 Fig**) (Krasileva *et al.*, 2017). The *ft-b2* mutations were verified using segregation analysis – mutant NILs were compared to *cv.* Cadenza and wild-type sibling lines.

Plants were grown at field sites of the John Innes Centre, Norwich, UK (52°62’25.7”N, 1°21’83.2”E) in 1 m^2^ plots, and in glasshouses under natural temperature and photoperiod conditions. Seeds were sown in week 2-3 of October. Phenotype data were collected over two growing seasons (2017 and 2018), and molecular data over two growing seasons (2018 and 2019). The *ft-b2.1* mutant and wild-type controls were grown in a glasshouse under 16 h light/ 8 h dark. The *ft-b2.2* and wild-type controls were grown under extra long-daylengths (22 h/ 2h).

For the moving bench experiment, *Ppd-D1a, ppd-1* and wild-type lines were grown in a glasshouse under natural photoperiods until the daylength reached 10 h. At 10 h, plants were shifted to a glasshouse with moving benches that transferred plants to a dark chamber after 10 h of natural daylight (**S7 Fig**). The experiment was repeated for two seasons.

### RNA extraction and expression analysis

Leaves from wild-type, *Ppd-D1a* and *ppd-1* NILs were sampled every 3-4 hours during the day and night, and at dusk (sunset). Sample time-points are expressed in terms of Zeitgeber time (ZT), with sunrise being 0 h. Leaf samples were harvested at photoperiods defined by 1 h increases in daylength, commencing at 9 h photoperiod and ending at 13 h (**Fig S1**). Three biological replicates were collected per time-point, each from the most recently emerged leaf of the primary tiller. For inflorescence expression analyses, samples were collected at four stages: vegetative (VG), double-ridge (DR), lemma primordia (LP) and terminal spikelet (TS). Each sample included pools of 5-15 inflorescences per replicate, dependent on stage. Three biological replicates were collected per stage.

Leaf RNA extractions were performed using the Spectrum Plant Total RNA Kit (Sigma-Aldrich). RNA extractions from developing inflorescences were performed using the RNeasy Plant Mini Kit (Qiagen). cDNA synthesis and RT-qPCR were performed as described previously, using a LightCycler480 Instrument II (Roche Life Science) (Boden *et al.*, 2014). RT-qPCR oligonucleotides are listed in Table S1 – the oligonucleotides amplify all three homoeoalleles for *FT1, FT2, VRN1* and the meristem identity genes, *Ppd-1* sequences are homoeoallele-specific. Candidate gene expression from leaf and inflorescence was normalized using TraesCS6D02G145100 (Traes_6DS_ BE8B5E56D.1), which we previously verified to be stably expressed in leaves and inflorescences across different photoperiods (Borrill *et al.*, 2016; Dixon *et al.*, 2018b). RT-qPCR data are the average of at least three biological replicates and two technical replicates per reaction.

The MADS-box transcription factors analysed here using the proposed nomenclature of (Schilling *et al.*, 2020), which are different from those we have described previously (Dixon *et al.*, 2018b).

### Inflorescence architecture measurements

Spikelet number was counted for the developing inflorescence at sequential stages. At the double-ridge stage, the spikelet meristem ridge was counted as a spikelet. From the lemma primordium stage onwards, spikelet meristems were clearly visible and counted as a spikelet. For fully emerged inflorescences, both viable and non-viable spikelets were counted. For *Ppd-D1a* insensitive *and ppd-1* lines, data of early inflorescence development are the average ± SEM of at least 3 replicates. For final spikelet numbers, data are the average ± SEM of at least 10 replicates. For *ft-b2* mutants, spikelet data are the average ± SEM of 5-7 replicates.

## RESULTS

### Seasonal and genetic regulation of *Photoperiod-1*

To investigate the seasonal regulation of the flowering pathway, we first measured the response of *Ppd-1* to increasing daylengths in the field. Specifically, we analysed expression of *Ppd-B1* and *Ppd-D1* in photoperiod-sensitive wild-type plants (*cv*. Paragon), as these homoeologues contribute the major photoperiod-insensitive alleles that confer early flowering phenotypes of hexaploid wheat in global breeding programmes (*Ppd-B1a* and *Ppd-D1a*, respectively) (Beales *et al.*, 2007; Diaz *et al.*, 2012; Shaw *et al.*, 2013). Transcripts were measured over a series of photoperiods defined by hourly increases in daylength from winter (9 h light/15 h dark) until late spring (13 h light/10 h dark), and diurnal patterns were analysed to precisely detect the daily peak(s) in gene expression (**Fig. 1; Fig. S1**). *Ppd-D1* and *Ppd-B1* were expressed at comparable levels to each other and they displayed very similar daily expression profiles (**Fig. 1A-B, D-E, Fig. S2**). The diurnal rhythm of *Ppd-D1* and *-B1* was maintained across all photoperiods, with transcripts peaking during the day (ZT 3-6 h) and at dusk, and dipping during the night (ZT 16-24 h). The consistent diel pattern in relation to dawn and dusk indicates *Ppd-1* expression adjusts to the changing daylengths. The amplitude of *Ppd-D1* expression was stable across all photoperiods tested, and *Ppd-B1* maintained a normalized expression range between 0.02 and 0.06, which was slightly higher at 10 and 13 h photoperiods (**Fig. 1A-B, D-E; Fig. S2**). These results suggest that the seasonal regulation of flowering in field-grown wheat is not determined by quantitative changes in *Ppd-1* expression.

**Figure 1:**
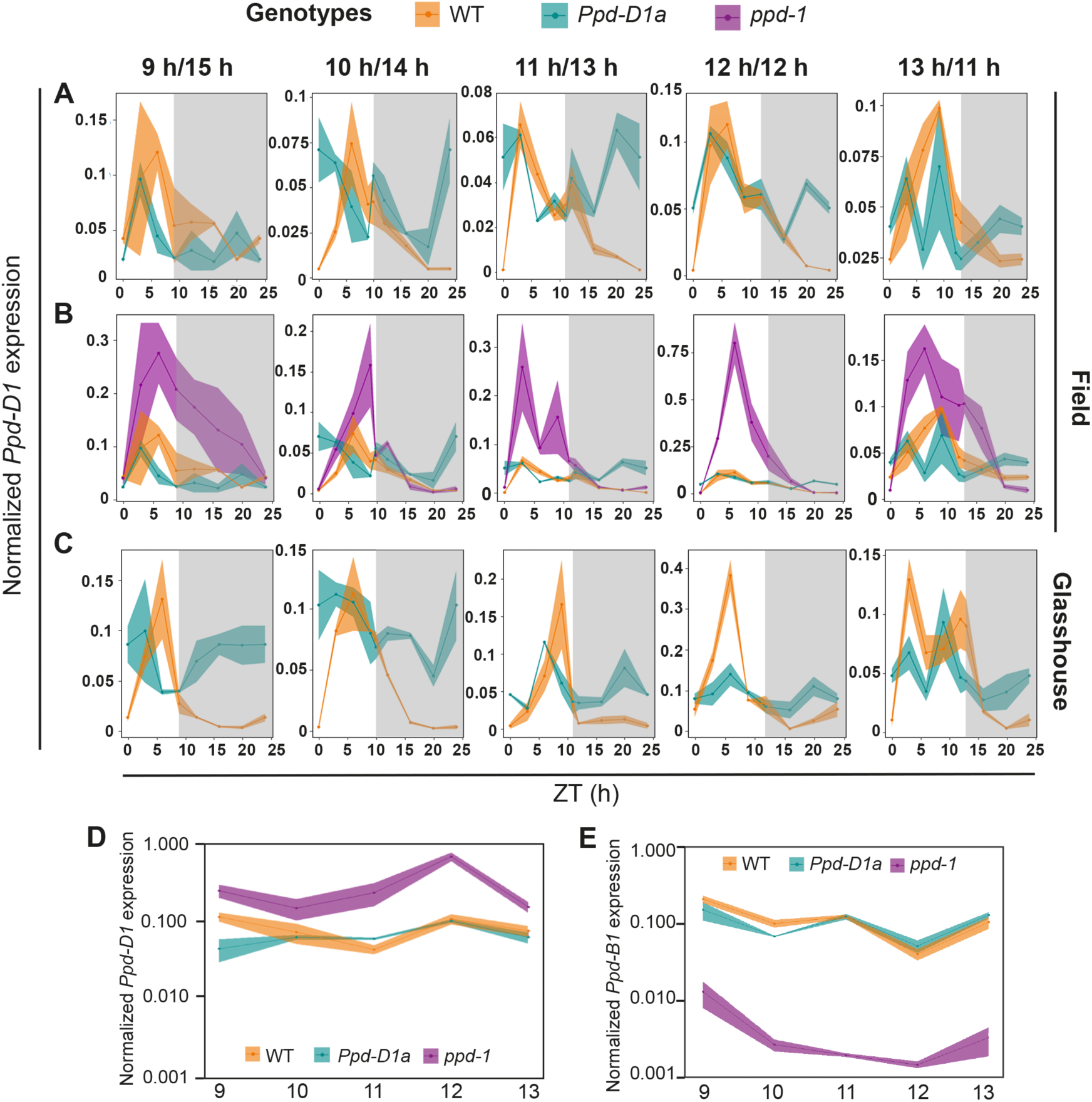
Seasonal regulation of *Ppd-1* under field- and glasshouse-based conditions. **(A, B)** Diurnal expression profiles of *Ppd-D1* in wild-type (orange), *Ppd-D1a* photoperiod-insensitive (cyan) and null *ppd-1* (magenta) NILs under field conditions. In **(A)**, the *ppd-1* data has been removed to show differences in transcript levels between wild-type and *Ppd-D1a* NILs. **(C)** Expression of *Ppd-D1* in WT and *Ppd-D1a* NILs under glasshouse conditions. **(D, E)** Data summarising the peak of **(D)** *Ppd-D1* and **(E)** *Ppd-B1* expression for each daylength, plotted on a logarithmic scale. The grey shading highlights night-time data points. All expression profiles are shown over a 24-hour period at hourly incremental increases in daylength, with time represented as Zeitgeber time, with sunrise being 0 h. Each point is the normalized mean transcript levels ± SEM of three biological replicates.

To determine how photoperiod-insensitive alleles modify *Ppd-1* expression under field conditions, we used a NIL expressing the early flowering *Ppd-D1a* allele that contains a 2.09 kb promoter deletion (Beales *et al.*, 2007; Shaw *et al.*, 2013). The photoperiod-insensitive *Ppd-D1a* allele altered the diurnal expression of *Ppd-D1* from the 10-13 h photoperiods by promoting higher expression late at night (ZT 20 and 24 h), relative to wild-type (**Fig. 1A**). This difference was particularly significant at 11 and 12 h daylengths; the difference in night-time expression was not detected in the 9 h photoperiod. There were also minor changes in expression of *Ppd-D1* during the day in the 13 h photoperiod (**Fig. 1A**). The insensitive *Ppd-D1a* allele did not significantly affect the amplitude of *Ppd-D1* expression, relative to wild-type, especially during the daytime when *Ppd-D1* peaked between 3-6 h after dawn in both genotypes (**Fig. 1A, D**). *Ppd-B1* expression was also affected in the photoperiod-insensitive NIL, particularly during the 10 and 11 h photoperiods, where it modified the timing of transcript peaks during the day (10 h) and increased the amplitude of *Ppd-B1* expression at certain time-points (**Fig. S2**). No significant effects on *Ppd-B1* expression were detected during the 12 and 13 h photoperiods. Based on these results, we conclude that the insensitive *Ppd-D1a* allele misregulates *Ppd-D1* expression during late hours of the night, particularly under day-neutral photoperiods of early spring. The altered expression of *Ppd-B1* in photoperiod-insensitive NILs suggests there is an interaction between *Ppd-1* homoeologues.

To genetically investigate the contribution of *Ppd-1* on seasonal regulation of flowering-time genes, we analysed NILs that contain non-functional alleles for all three homeoalleles (i.e. *ppd-1* NILs). These lines carry deletions for *Ppd-A1* and *Ppd-B1* and a nonsense allele of *Ppd-D1*, which produces a transcript with a premature stop-codon (Shaw *et al.*, 2013). *Ppd-D1* transcript levels were significantly higher during the day of the 10-13 h photoperiods, relative to wild-type, and, as expected, no *Ppd-B1* transcripts were detected (**Fig. 1B, D, Fig. S3**). These results indicate that *Ppd-1* may form part of a self-regulatory feedback loop that represses *Ppd-D1* expression during the day, which is disrupted when there is no functional *Ppd-1* protein.

An unexpected outcome of the field-based *Ppd-D1* expression analysis was that misregulated expression of *Ppd-D1* in the photoperiod-insensitive line was limited to late hours of the night of the 10-12 h photoperiods, as previous analyses using controlled environments showed insensitive *Ppd-D1a* alleles alter expression during all hours of the evening, particularly under short-day photoperiods (9 h/15 h) (**Fig. 1A**) (Beales *et al.*, 2007; Shaw *et al.*, 2012; Boden *et al.*, 2015). To investigate whether there is a difference between controlled-environment and field conditions, we compared the expression of *Ppd-D1* from field- and glasshouse-grown plants (**Fig. 1C; Fig. S3**). We found that the photoperiod-insensitive allele promoted significantly higher expression of *Ppd-D1* during all hours of the night in all photoperiods for glasshouse-grown plants, relative to wild-type, consistent with previous reports that used controlled environments (**Fig. 1C**) (Beales *et al.*, 2007; Shaw *et al.*, 2012; Boden *et al.*, 2015). These data indicate that controlled growth conditions exaggerate the impact of photoperiod-insensitive alleles on *Ppd-D1* expression, relative to field-grown plants. In wild-type, the differences in *Ppd-D1* expression between field- and glasshouse-grown plants were less dramatic – the transcript peaks were slightly higher in glasshouse-grown plants during the day and the troughs were moderately lower during the night, relative to the field (**Fig. 1C**). In the *ppd-1* NIL, we observed a similar dramatic increase in *Ppd-D1* expression that was detected in the field, supporting the suggestion that *Ppd-1* forms a self-regulatory feedback loop (**Fig. S3**).

### Step-wise increases in *FLOWERING LOCUS T1* expression underpin seasonal regulation of flowering

To investigate the seasonal progression of major flowering-time genes that act downstream of *Ppd-1*, we analysed expression of *FT1* and *VERNALIZATION1* (*VRN1*) (**Fig. 2, Fig. S4**). In wild-type, *FT1* expression was very low under 9 and 10 h daylengths and moderately induced between 10 and 11 h photoperiods of late winter, with a 3-fold increase in transcripts (**Fig. 2A-B, D**). Under these daylengths, the diurnal expression pattern of *FT1* included peaks during the day and at dusk. This pattern continued under 12 h photoperiods with a significant peak in the morning (Zeitgeber Time (ZT) 3 h) and a minor peak after dusk (ZT 16 h). *FT1* transcripts significantly increased in spring between 12 and 13 h daylengths, with expression at 13 h photoperiods being approximately 10-fold higher than that of 12 h (**Fig. 2A, D**). Under 13 h daylengths, *FT1* transcripts peaked during the day at ZT 6 h and again in the evening (ZT 13-16 h). The *ppd-1* lines unexpectedly displayed a similar diurnal pattern of *FT1* expression and near-identical responsiveness to the changes in daylength, relative to wild-type; however, the amplitude of expression was significantly lower than wild-type at all time-points of the 11-13 h photoperiods (**Fig. 2A, D**). Conversely, the insensitive *Ppd-D1a* plants showed dramatically different *FT1* transcript activity compared to wild-type, with expression detected much earlier than wild-type during the 9-10 h photoperiods and showing much higher amplitudes (**Fig. 2B, D**). In the *Ppd-D1a* NILs, we detected low levels of *FT1* transcripts under 9 and 10 h photoperiods of winter, before expression increased significantly as daylengths extended at the beginning of spring to 11 h, with levels comparable to that detected in wild-type under 13 h (**Fig. 2B, D**). *FT1* expression then settled at 12 h daylengths, before increasing again at 13 h – under photoperiods of 12 and 13 h, the diurnal expression pattern included peaks during the morning (ZT 3-6 h) and evening (ZT 16-20 h). These results show that *Ppd-1* is required for robust expression of *FT1*, and that photoperiod-insensitive alleles promote rapid induction of *FT1* under shorter daylengths of winter. In wild-type, the increase in *FT1* expression occurs in a step-wise process, with an initial rise at the beginning of spring (11 h) followed by a stronger induction in late-spring (13 h; **Fig. 2D**). The relative differences in *FT1* expression associated strongly with flowering-time phenotypes, as the photoperiod-insensitive lines flowered 13 days earlier than wild-type, while the *ppd-1* NILs flowered 11 days later (**Fig. 2E**).

**Figure 2:**
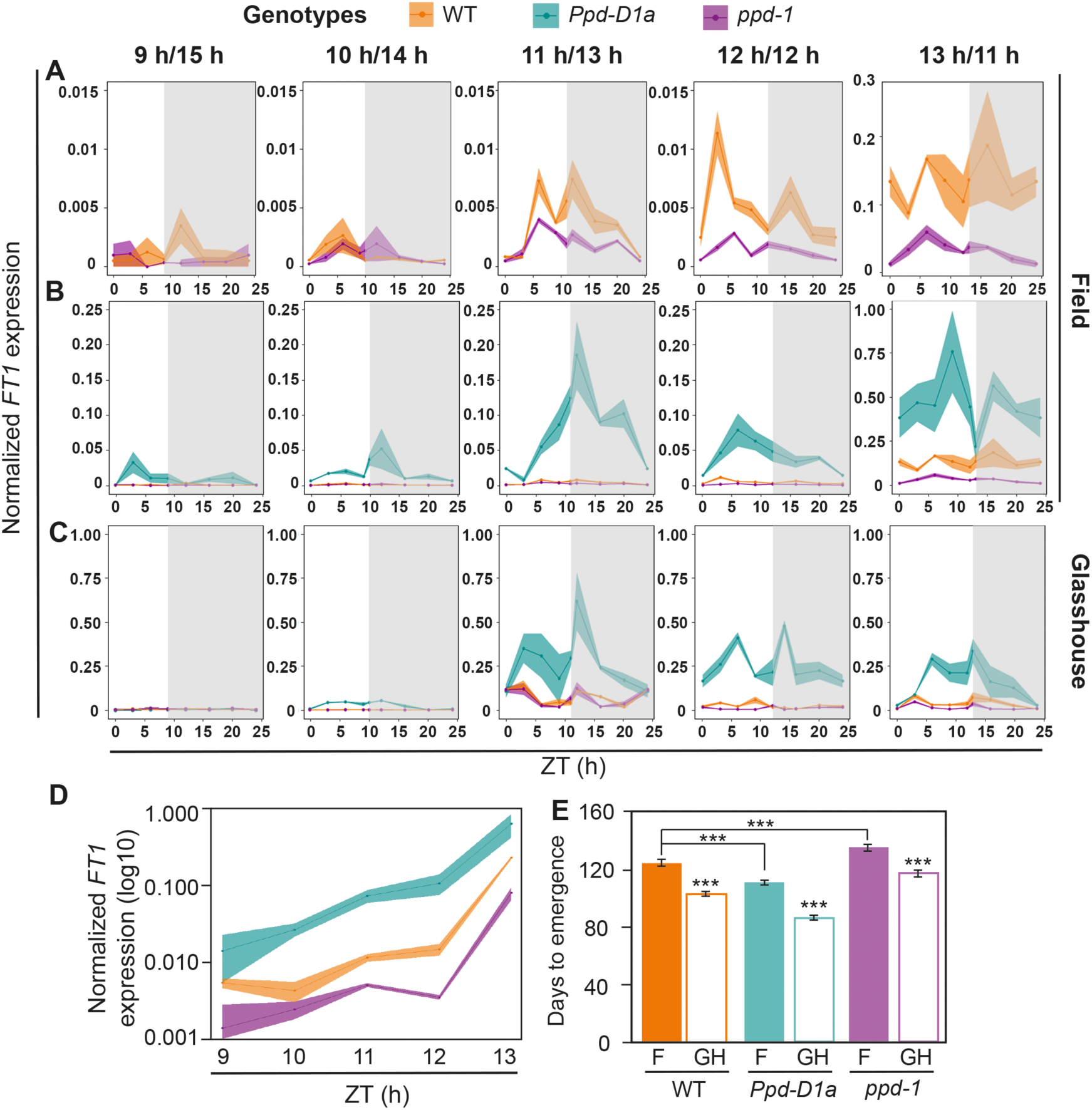
Seasonal regulation of *FT1* under field- and glasshouse-based conditions. Diurnal expression profile of *FT1* in wild-type (orange), *Ppd-D1a* photoperiod-insensitive (cyan) and null *ppd-1* (magenta) NILs under field **(A, B)** and glasshouse **(C)** conditions. In **(A)**, the data from the *Ppd-D1a* NIL has been removed to show differences in transcript levels between wild-type and *ppd-1* NILs. **(D)** Data summarising the peak of *FT1* expression for each daylength, plotted on a logarithmic scale. **(E)** Field and glasshouse flowering time phenotypes for the three *Ppd-1* NILs, normalized as days to emergence from the 9 h daylength. The grey shading highlights night-time data points. **(A-D)** Each point is the normalized mean transcript levels ± SEM of three biological replicates, **(E)** data is mean ± SEM, with field replicates being 5 independent plots and glasshouse replicates being 10-15 plants. *** *P <* 0.001.

As we detected a difference in *Ppd-1* expression in field-grown plants relative to glasshouse conditions, we hypothesised that *FT1* would display an altered transcriptional profile under the two growth regimes. Contrary to the field, wild-type glasshouse-grown plants expressed *FT1* at significant levels under the 11 h photoperiod, which stabilized as daylengths increased to 12 and 13 hours, indicating that 11 h daylengths are sufficient to induce flowering in wheat (**Fig. 2C**). The *ppd-1* lines displayed a similar trend with considerable expression at 11 h, which is maintained through to 13 h; however, at 12 and 13 h there are significantly fewer transcripts than wild-type throughout the day and night (**Fig. 2C**). In the photoperiod-insensitive lines, *FT1* was induced at the 10 h daylength, with a dramatic increase in expression as daylengths extend to 11 h, which is maintained through to 13 h (**Fig.2C**). At all time-points and daylengths, the amplitude of *FT1* transcripts is higher in the *Ppd-D1a* lines than wild-type and *ppd-1* NILs – *FT1* transcript levels of photoperiod-insensitive plants grown at 11 h are comparable to those of wild-type plants at 13 h (**Fig. 2C**). A similar diurnal expression pattern of *FT1* was detected in all three lines, with one peak detected in the morning (ZT 0-6 h) and another at dusk. The relative differences in *FT1* expression are reflected in the flowering-time phenotypes, as photoperiod-insensitive lines flowered 20 days earlier than wild-type, while the *ppd-1* NILs flowered 11 days later (**Fig. 2E**).

To investigate floral promoting genes that respond to temperature, we examined the transcript levels of *VRN1*, whose expression increases following exposure to cold and influences the timing of flowering for winter wheat treated by warm temperatures (Danyluk *et al.*, 2003; Trevaskis *et al.*, 2003; Yan *et al.*, 2003; Dixon *et al.*, 2019) (**Fig. S4**). *VRN1* was strongly expressed in leaves under all photoperiods. While *VRN1* did not display a particular pattern of diurnal expression across all photoperiods, transcript levels were significantly affected by *Ppd-1* at certain time-points of the day and night. *VRN1* expression was significantly higher in photoperiod-insensitive NILs, relative to wild-type, and significantly lower in *ppd-1* NILs (**Fig. S4**). These data indicate the early flowering phenotype of photoperiod-insensitive wheat involves a coordinated increase in expression of floral activating genes, including *VRN1* and *FT1*.

### Inflorescence development responds dynamically to changes in FT1 activity

To investigate the connection between the floral promoting pathway in the leaves with IM development, we first examined developmental progression of inflorescences from field-grown plants in the context of changes in *FT1* expression (**Fig. 3**). In wild-type, the SAM remained vegetative until the beginning of spring at the 11 h photoperiod, at which point it transitioned towards the double-ridge stage (**Fig. 3A-C, Fig. S5**). The timing of this transition coincided with the three-five-fold increase in *FT1* expression that occurs between the 10 and 11 h photoperiods (**Fig. 2A, D**). The IM remained at the double-ridge stage until daylengths extended to 12.5 h, when it transitioned to the glume and lemma primordium stages (**Fig. 3A; Fig. S5**). The timing of this second transition coincided with the ten-fold increase in *FT1* expression that occurs in leaves between the 12 and 13 h photoperiods of April (**Fig. 2A, D**). The IMs remained at the lemma primordium stage until daylengths reached 13.5 h. (**Fig. 3A, Fig. S5**). Growth and development of the IM then proceeded rapidly beyond this point, with spikelets at the lemma primordium stage forming floret primordia and reaching the terminal spikelet stage when daylengths were 14 h (**Fig. 3A, Fig. S5**). Interestingly, transition of the wild-type IM from glasshouse-grown plants followed a very similar progression, only that key events occurred at a photoperiod approximately one hour prior, relative to the field, to coincide with the earlier induction of *FT1* expression (**Fig. 3B**). The IMs transitioned to double-ridge and lemma primordium stages when daylengths were 10 and 11.75 h, respectively (**Fig. 3B**). These results show that the timing of the transition for wild-type IMs to the double-ridge stage in the field coincides with an initial rise in *FT1* expression. The IMs then stall at the double-ridge stage until the second increase in *FT1* expression, when the IMs proceed to the lemma primordium and terminal spikelet stages. In the *Ppd-D1a* insensitive NILs, IM development closely tracked changes in *FT1* expression. The IM transitioned to the double-ridge stage between the 10-11 h daylengths of late winter, coinciding with the photoperiods when *FT1* transcripts increased significantly (**Fig. 3A, Fig. S5**). The IMs then proceeded rapidly to the glume and lemma primordium stages when daylengths were 11 and 12 h, respectively, without the developmental pause observed in wild-type, which coincided with the high expression of *FT1* under these photoperiods (**Fig. 2, 3A**). The IMs arrived at the terminal spikelet stage when daylengths were 12.75 h. The IMs of *ppd-1* NILs transitioned to the double-ridge stage at the 11 h photoperiod, proceeding to the lemma primordium and terminal spikelet stages when daylengths were 14 and 15 h, respectively (**Fig. 3A, Fig. S5**). Development of the *ppd-1* inflorescences therefore closely followed changes in *FT1* expression in leaves, which initiated between the 10 and 11 h photoperiods, before increasing again at 13 h. In both the *Ppd-D1a* photperiod-insensitive and *ppd-1* NILs, the influence of *Ppd-1* on spikelet number was detected between glume primordium and terminal spikelet stages, with fewer spikelets forming in the insensitive NIL and more developing in *ppd-1*, relative to wild-type (**Fig. 3C**). This trend in spikelet number for each genotype was also observed at maturity, with photoperiod-insensitive lines producing shorter inflorescences with fewer spikelets than wild-type, while *ppd-1* NILs formed significantly longer inflorescences with more spikelets (**Fig. S6**). Taken together with the analysis of *FT1* transcript levels, these results show that developmental progression of the IM is dynamically linked with changes in *FT1* expression in the leaves, and variant alleles of *Ppd-1* modify the timing and rate of inflorescence development to alter spikelet number.

Our previous analysis showed *FT1* is required for robust expression of meristem identity genes and timely progression to the terminal spikelet stage (Boden *et al.*, 2015; Dixon *et al.*, 2018a). Based on the *FT1* expression analyses and modified development of IMs in *Ppd-D1a* and *ppd-1* NILs shown here, we hypothesised that expression of meristem identity genes occurs earlier and to a higher level in insensitive NILs, relative to wild-type, and is delayed and lower in the *ppd-1* NILs. To test this, we analysed expression of MADS-box transcription factors that have key roles in early inflorescence development, including the regulation of spikelet architecture, floral organ identity and fertility that are affected by *Ppd-1* allelism (Boden *et al.*, 2015; Schilling *et al.*, 2020). The genes include *APETALA1-like* (*AP1-2, AP1-3* and *VRN1/AP1-1*), *AGAMOUS-LIKE6* (*AGL6*), *SEPALLATA1-6 (SEP1-6)* and *SUPPRESSOR OF CONSTANS1* (*SOC1*), as well as *GRAIN NUMBER INCREASE1* (*GNI1*) and *FLOWERING LOCUS T2* (*FT2*) that regulate floret fertility and spikelet number in wheat (**Fig. 4**) (Boden *et al.*, 2015; Sakuma *et al.*, 2019; Shaw *et al.*, 2019; Schilling *et al.*, 2020). Transcripts of *AP1-2, AP1-3, SEP1-6* and *SOC1* were detected in *Ppd-D1a* NILs earlier than wild-type, as determined by time since germination (daylength) (**Fig. 4A**). These genes were expressed between the 10-11 h photoperiods in *Ppd-D1a* NILs, but not in wild-type until the 12-13 h photoperiods, coinciding with daylengths of mid-spring when *FT1* expression increased significantly in leaves and the IMs transitioned to the double-ridge stage. Expression of *FT2, AGL6* and *GNI1* occurred later in the season with transcripts detected earlier in *Ppd-D1a* NILs, relative to wild-type. For the *ppd-1* NILs, expression of these genes was inverted relative to changes observed in the insensitive NILs, with transcripts detected at later photoperiods than wild-type, at daylengths corresponding to the delay in *FT1* induction and IM development observed in these plants. The amplitude of *AP1-2, AP1-3, VRN1, SEP1-6, SOC1, MADS6* and *FT2* transcripts unexpectedly reached the same maximum level in wild-type as in *Ppd-D1a* NILs, demonstrating that meristem identity genes are not expressed at higher levels in photoperiod-insensitive plants (**Fig. 4**). Conversely, transcripts were significantly lower in the *ppd-1* NILs, showing *Ppd-1* is required for robust expression of genes that promote spikelet and floret development (**Fig. 4**). *GNI1* transcripts spiked to higher levels in *Ppd-D1a* NILs, relative to wild-type and *ppd-1*, at the green anther stage. Based on the seasonal shift in expression peak detected for these genes in the insensitive and *ppd-1* NILs, relative to wild-type, we hypothesised that their expression was changing in relation to the developmental stage rather than daylength. To test this hypothesis, we normalised gene expression according to developmental stages including vegetative, double-ridge, lemma primordium, terminal spikelet, white anther and green anther (**Fig. 4B**). Following normalization, transcript levels were almost identical in *Ppd-D1a* NILs, relative to wild-type, supporting the conclusion that photoperiod-insensitive alleles advance expression of meristem identity genes without increasing transcript levels (**Fig. 4B**). An exception to this trend was *FT2*, which was higher in *Ppd-D1a* NILs at the lemma primordium stage, relative to wild-type, in which transcript levels did not increase significantly until terminal spikelet (**Fig. 4B**). In *ppd-1* NILs, transcripts of meristem identity genes were much lower and peaked at later developmental stages, relative to wild-type (**Fig. 4B**). These field-based results show insensitive alleles provoke accelerated but equal expression of meristem identity genes in IMs, relative to wild-type, and that *Ppd-1* is required for timely and robust induction of genes that promote spikelet and floret development. The expression of meristem identity genes for all three genotypes is dynamically linked with the activity of *FT1* in the leaves.

**Figure 3:**
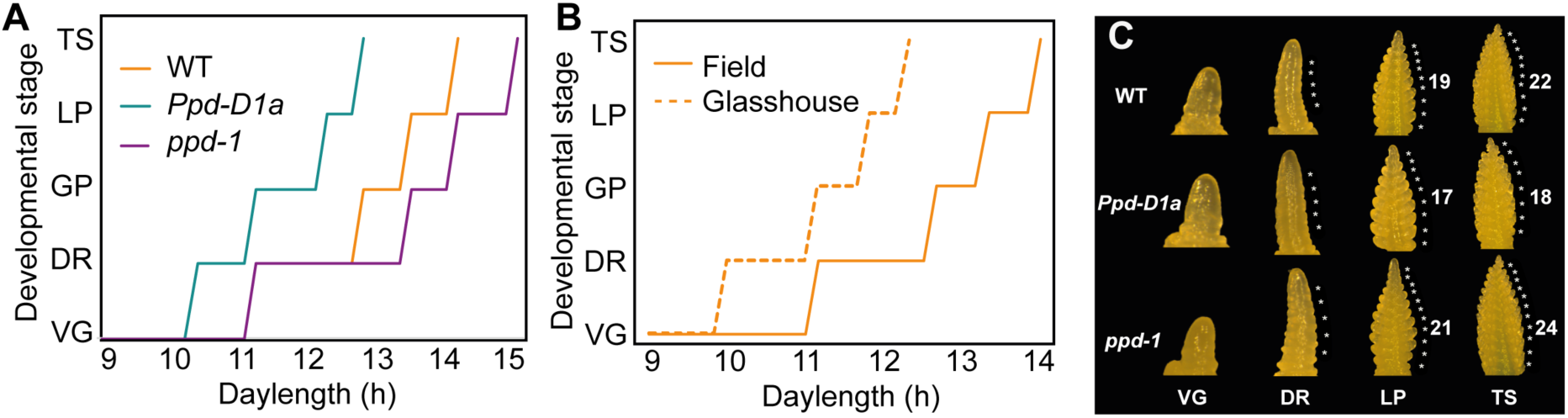
Seasonal progression of inflorescence meristem development in *Ppd-1* NILs. **(A)** Progression of inflorescence meristem development throughout the season in wild-type (orange), *Ppd-D1a* photoperiod-insensitive (cyan) and null *ppd-1* (magenta) NILs under field conditions, at the vegetative (VG), double ridge (DR), glume primordia (GP), lemma primordia (LP) and terminal spikelet (TS) stages. **(B)** Comparative progression of wild-type inflorescence meristems grown under field (solid) and glasshouse (dashed) conditions. (**C**) Representative images of inflorescence meristems from each genotype at the four developmental stages. **(A-B)** Data are the average of 4-5 replicates per developmental stage.

**Figure 4:**
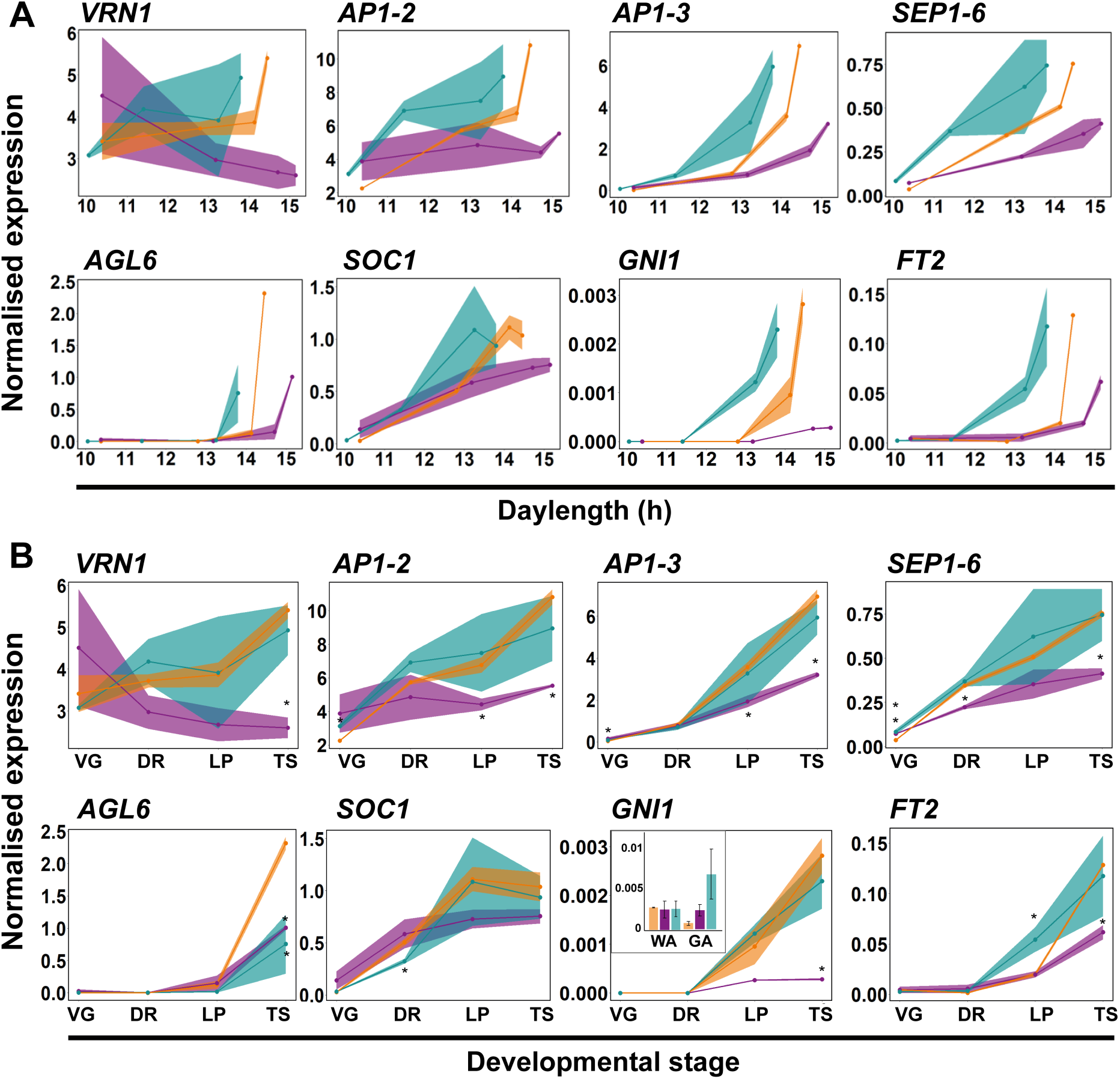
Seasonal and stage-specific expression analysis of meristem identity genes. Expression of key meristem identity genes plotted by daylength **(A)** and developmental stage **(B)** from plants grown under field conditions. Data includes expression profiles for wild-type (orange), *Ppd-D1a* photoperiod-insensitive (cyan) and null *ppd-1* (magenta) NILs. The defined stages are vegetative (VG), double ridge (DR), lemma primordium (LP), terminal spikelet (TS), white anther (WA) and green anther (GA). Data are the mean ± SEM of three-four biological replicates.. * *P <* 0.05

### *FT2* contributes to the termination of spikelet development

To further investigate the role of the initial induction of *FT1* expression in leaves and its relationship to IM development, we analysed the molecular and physiological effects of maintaining plants at 10 h daylengths of late winter. We hypothesised that the inceptive rise in *FT1* expression in wild-type promotes the initial stages of inflorescence development, and the second stronger induction is required to proceed to later reproductive stages. To test this hypothesis, plants were grown in a glasshouse under natural photoperiods until the daylength was 10 h, before being shifted to a moving bench that maintained a fixed short-day 10 h photoperiod (**Fig. S7**). Wild-type plants maintained at 10 h progressed to the lemma primordium stage on the same day and produced the same amount of spikelets as plants maintained under natural photoperiods, by which time the daylength had surpassed 12 h (**Fig. 5A**). However, these plants stalled at the lemma primordium stage for an extra 30 days before arriving at the terminal spikelet stage and produced more spikelets compared to plants maintained under natural photoperiods (24 ± 0.3 vs 29 ± 0.5 spikelets, respectively; **Fig. 5A-C**). The delay in inflorescence development coincided with the developmental stage at which the second stronger induction of *FT1* expression occurred (**Fig. 2A, D**). This delay was also observed in *ppd-1* NILs, which produced more spikelets and transitioned to the terminal spikelet stage significantly later than plants maintained under natural photoperiods (25.4 ± 0.2 vs 28.3 ± 0.3 spikelets; **Fig. 5B-C**). The *Ppd-D1a* NILs transitioned to the terminal spikelet stage more rapidly than wild-type, but there was a slight delay relative to natural photoperiods, as plants maintained at 10 h produced more spikelets than glasshouse-grown plants (21.5 ± 0.2 vs 25.8 ± 0.4; **Fig. 5B-C**). Wild-type and *ppd-1* NILs flowered later under the fixed 10 h photoperiods than plants grown under natural daylengths, but the photoperiod-insensitive NILs flowered at the same time (**Fig. S7**). In addition to increasing spikelet number, the 10 h photoperiods altered inflorescence architecture of wild-type and the *ppd-1* NIL by forming elongated basal internodes immediately prior to ear emergence, as occurs in *ppd-1* mutants grown under constant long-days (**Fig. 5D; Fig. S7**) (Shaw *et al.*, 2013; Boden *et al.*, 2015). The insensitive *Ppd-D1a* lines did not produce these elongated internodes. These results indicate that the initial induction of *FT1* is sufficient to promote transition of the IM to the lemma primordium stage; however, the second higher induction of *FT1* is required to progress development of the IM to the terminal spikelet and later stages. This delay is overridden in *Ppd-D1a* insensitive NILs, most likely due to the higher expression of *FT1* (**Fig. 2**).

**Figure 5:**
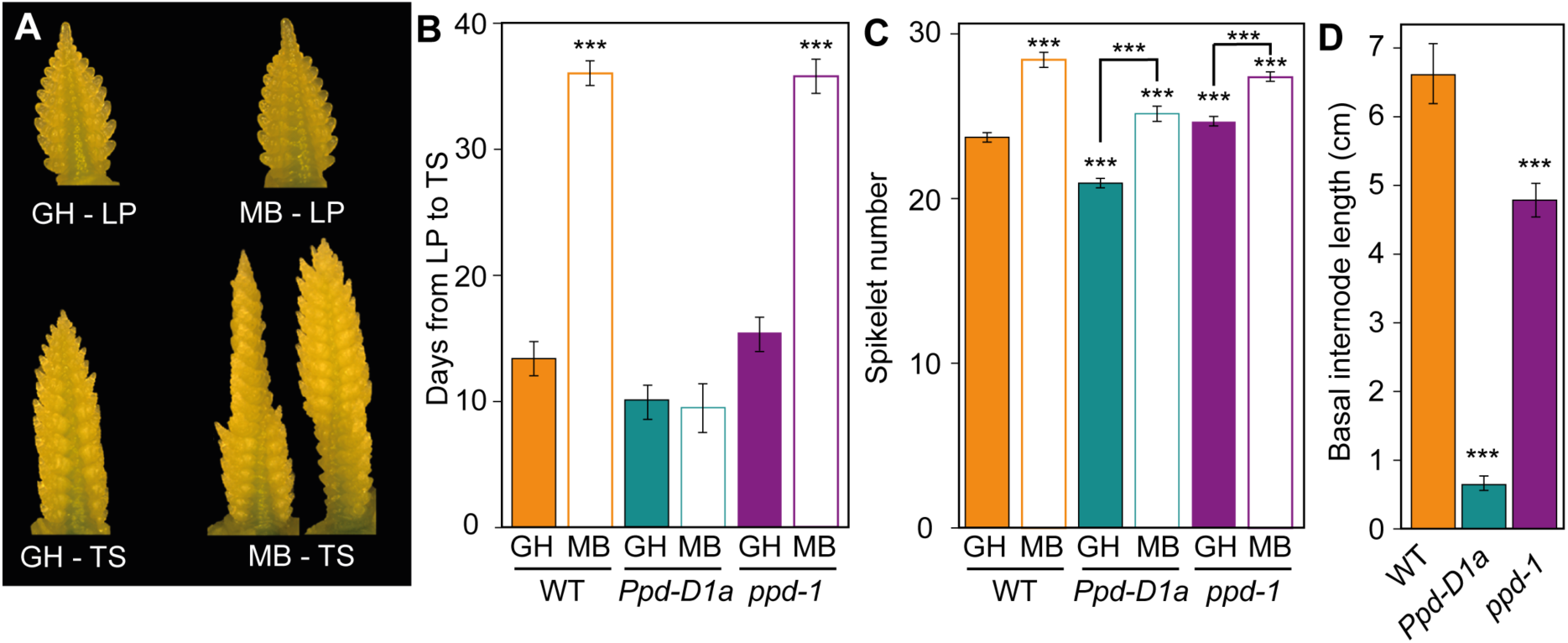
Inflorescence phenotypes of plants maintained under short-day photoperiods. **A)** Inflorescences of wild-type plants grown under natural glasshouse (GH) and 10 h moving bench (MB) conditions develop at the same rate until the lemma primordia (LP) stage, but develop more spikelets by the terminal spikelet stage (TS). **(B)** IM development of WT and *ppd-1* NIL plants grown under MB conditions is significantly delayed between the LP and TS stages. **(C)** Spikelet numbers for all three genotypes grown under natural GH or MB conditions. **(D)** Wild-type and *ppd-1* NIL plants show elongated basal rachis internodes, but not *Ppd-D1a* NILs. Data are the average ± SEM of five biological replicates. *** *P <* 0.001.

To investigate genes that contribute to the progression of inflorescence development beyond the lemma primordium stage, we investigated the role of *FT2*. We selected *FT2* because its expression increased dramatically in wild-type inflorescences between the lemma primordium and terminal spikelet stages, and it was expressed earlier in *Ppd-D1a* NILs and later in *ppd-1* plants (**Fig. 4**). Expression of *FT2* was much lower in IMs at the glume primordium stage of plants shifted to the 10 h photoperiod, relative to plants maintained in natural photoperiods, suggesting development of the IM to the terminal spikelet stage is associated with robust expression of *FT2* (**Fig. 6A**). To test the role of *FT2* genetically, we analysed two independent lines containing missense mutations in *FT2* of the B genome (*FT-B2*; G45D and R150C), which are predicted to be deleterious for protein function (PROVEAN scores of -5.1 and -6.8) (Choi & Chan, 2015). The *ft-b2* mutants produced more spikelets than their wild-type NILs, indicating progression of the IM to the terminal spikelet stage was delayed (**Fig. 6B-D**). The *ft-b2* mutants flowered later than wild-type, even under extreme long-day photoperiods (**Fig. 6E**). These results indicate *FT2* has an important role in determining the completion of spikelet development, coinciding with the second induction of *FT1* expression.

**Figure 6:**
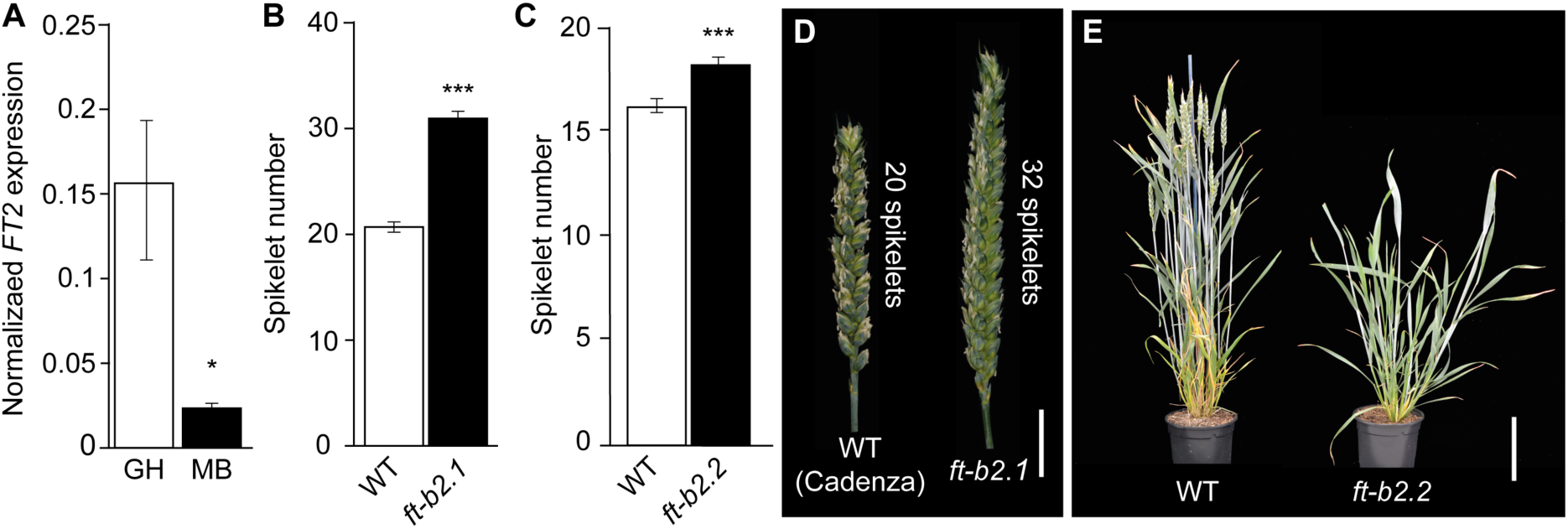
*FT2* influences inflorescence development and spikelet number. **(A)** *FT2* expression is perturbed in developing IMs of wild-type plants shifted to 10 h photoperiods using a moving bench (MB), relative to plants maintained under natural photoperiods. **(B-D)** Spikelet number increases on inflorescences of (**B**) *ft-b2.1* mutants and (**C**) *ft-b2.2* mutants, relative to wild-type, (**D**) including images of representative inflorescences. (**E**) *ft-b2.2* mutant lines flower later that wild-type (WT) under long-daylengths. **(A)** Data are the mean ± SEM of three biological replicates, and five replicates for **(B-C).** * *P <* 0.05; *** *P <* 0.001.

## DISCUSSION

Flowering is a crucial process in the lifecycle of an annual plant. Flowering-time genes respond to seasonal cues including daylength and temperature to coordinate seed production with favourable environmental conditions. Our understanding about the genes that regulate flowering of wheat mostly stem from work performed under controlled conditions different from those experienced by field-grown plants (Worland *et al.*, 1998; González *et al.*, 2005; Beales *et al.*, 2007; Boden *et al.*, 2015; González-Navarro *et al.*, 2015; Prieto *et al.*, 2018; Perez-Gianmarco *et al.*, 2019). Here, we provide new knowledge about the seasonal regulation of flowering using natural photoperiods of a standard growing season.

The induction of *FT-like* genes is core to the floral transition of angiosperms, which is conserved in wheat with *FT1* being a key activator of flowering (Yan *et al.*, 2006; Bratzel & Turck, 2015; Dixon *et al.*, 2018a; Finnegan *et al.*, 2018). Our analysis of field-grown plants unexpectedly showed that *FT1* induction occurs in a step-wise process, with an initial rise in transcripts detected under 11 h daylengths before increasing again as they extend to 13 h (**Fig. 7**). The induction of *FT1* under 11 h photoperiods demonstrates that long daylengths are not necessary to induce *FT1* expression in photoperiod-sensitive wheat. This is consistent with inflorescences transitioning to early reproductive stages at the beginning of spring, when daylengths are shorter than 12 h – a process promoted by *FT1* (González-Navarro *et al.*, 2015; Dixon *et al.*, 2018a). The rise in *FT1* expression as daylengths extend to 13 h is consistent with the significant increase in *FT1* transcripts when plants are shifted to extreme long-days (22 h/2 h), and with analysis of Arabidopsis grown under natural summer photoperiods, where *FT1* transcripts were higher under 16 h than 14 h photoperiods (Dixon *et al.*, 2018b; Song *et al.*, 2018). A second unexpected outcome of the *FT1* expression analysis regards the diurnal expression pattern under natural photoperiods, which showed peaks in transcript levels in the morning and at dusk. These patterns are different from those reported for wheat and barley in controlled conditions under constant long-days, where *FT1* peaks at dusk (Turner *et al.*, 2005; Campoli *et al.*, 2012; Chen *et al.*, 2014; Boden *et al.*, 2015). Our data is consistent with comparisons of Arabidopsis grown under laboratory versus natural photoperiod conditions, which showed temperature and light quality signals are responsible for a predominant peak of *FT* expression during the morning that was not prevalent in the laboratory (Song *et al.*, 2018). These data show that the regulation of flowering-time processes in the field are potentially more complex than indicated by work performed using controlled conditions.

**Figure 7:**
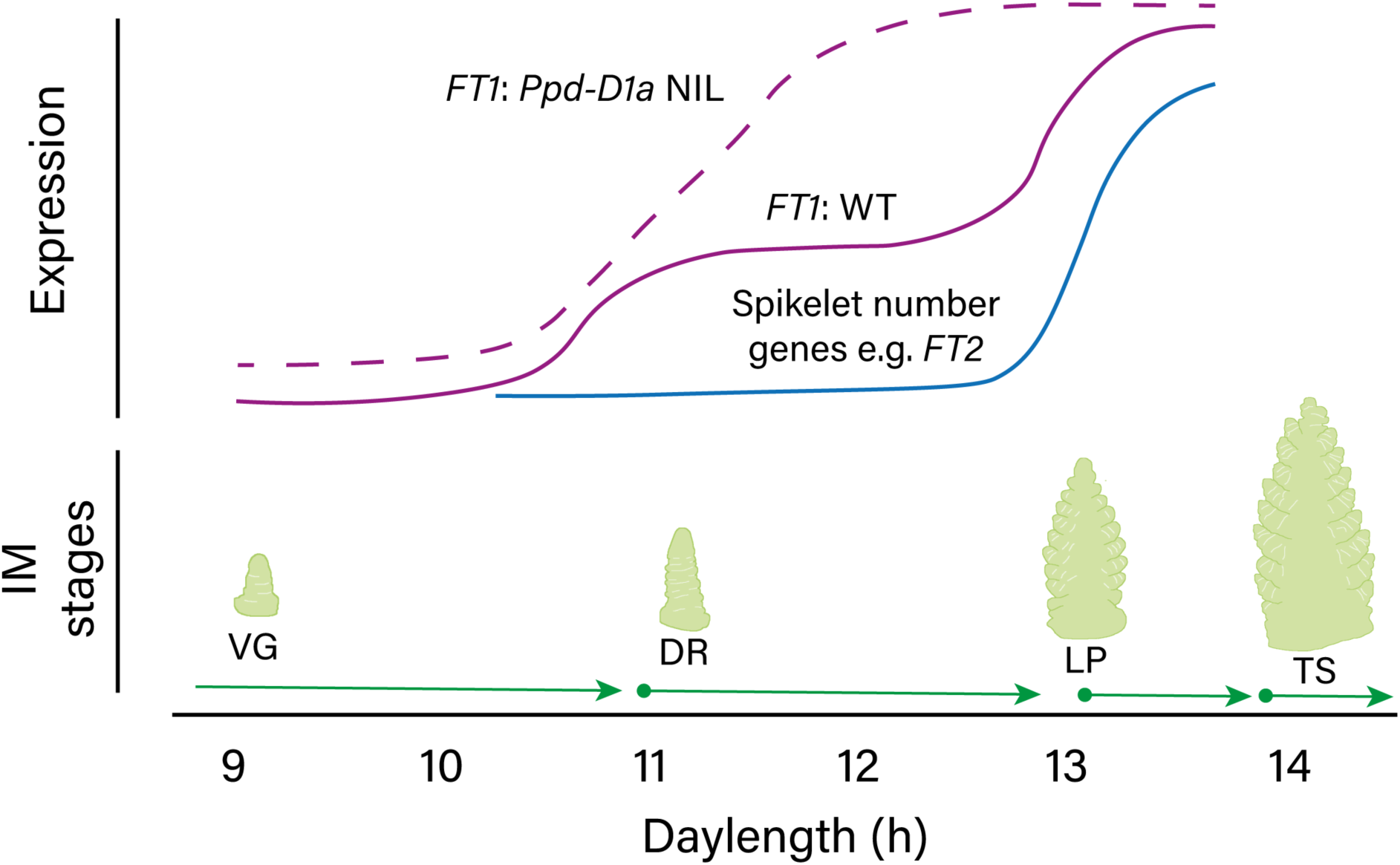
Model for seasonal regulation of flowering-time pathways and inflorescence development in bread wheat. A model outlining the seasonal regulation of *FT1* expression in wheat, defined by hourly increases in daylength, in wild-type (solid, purple line) and *Ppd-D1a* photoperiod-insensitive lines (dashed purple line). The second induction of *FT1* promotes expression of meristem identity genes that determine spikelet number, such as *FT2* (blue), which helps transition the IM to the lemma primordium and terminal spikelet stages. The regulation of floral promoting pathways coincides with developmental stages (cartoon images) according to the seasonal progression of inflorescence development, shown here for wild-type plants. The stages include vegetative (VG), double ridge (DR), lemma primordium (LP) and terminal spikelet (TS).

Genetic variation for *Ppd-1* has a major effect on *FT1* activity and flowering time, with photoperiod-insensitive alleles promoting higher *FT1* expression and earlier flowering, relative to wild-type (Beales *et al.*, 2007; Wilhelm *et al.*, 2009; Shaw *et al.*, 2012; Shaw *et al.*, 2013; Boden *et al.*, 2015). Here, we have shown that photoperiod-insensitive alleles cause *FT1* to be expressed earlier in the season and to higher levels, relative to wild-type, with the initial levels comparable to those detected in wild-type later in the year (**Fig. 7**). The photoperiod-insensitive allele significantly affects *FT1* expression during early spring, which corresponds to photoperiods when inflorescence development initiates in the field and spikelet number is determined (González-Navarro *et al.*, 2015). The pronounced effect of the *Ppd-D1a* allele on *FT1* expression during shorter daylengths of winter is consistent with the exaggerated acceleration of flowering that occurs in photoperiod-insensitive lines under constant 9 h photoperiods, where flowering occurs 60-190 days earlier than photoperiod-sensitive lines; photoperiod-insensitive alleles only accelerate flowering by 6-12 days under field conditions (**Fig. 2E**) (Worland *et al.*, 1998; Beales *et al.*, 2007; Diaz *et al.*, 2012; Cane *et al.*, 2013). Interestingly, the misregulated expression of *Ppd-D1* in photoperiod-insensitive lines was less dramatic in field-grown plants, relative to the glasshouse and laboratory conditions, with the high evening expression not detected under 9 h photoperiods and limited to late hours of the night as photoperiods extended (Beales *et al.*, 2007; Boden *et al.*, 2015). Given the glasshouse- and field-grown plants were both grown under natural photoperiods, the altered pattern of *Ppd-1* expression may be attributed to differences in temperature and/or light quality, which would be consistent with *Ppd-1* being regulated by phytochromes that are sensors for changes in light quality and temperature in plants (Rockwell *et al.*, 2006; Chen *et al.*, 2014; Jung *et al.*, 2016). Regarding the *ppd-1* lines, *FT1* transcript levels were significantly lower than wild-type, but *FT1* expression still responded to increasing daylengths and displayed the same diurnal pattern as wild-type. These results indicate that *Ppd-1* is not the only factor regulating seasonal activity of *FT1*.

In addition to a role regulating *FT1*, the higher levels of *Ppd-D1* transcripts in *ppd-1* lines indicates that *Ppd-1* may influence its own expression through a self-regulatory feedback loop. This potential function of *Ppd-1* is supported by the interaction detected between *Ppd-1* homoeologues in the photoperiod-insensitive *Ppd-D1a* NILs, and is consistent with *Ppd-B1* expression being higher in *Ppd-D1a* mutants containing splice site mutations (Boden *et al.*, 2015).

The process of flowering involves communication of FT1 protein from the leaves to the SAM, from which reproductive development is promoted through expression of meristem identity genes (Shaw *et al.*, 2013). In Arabidopsis and rice, overexpression of *FT*/*Hd3a* (*Heading date 3a*) leads to hyper-activation of meristem identity genes (Kardailsky *et al.*, 1999; Yoo *et al.*, 2005; Taoka *et al.*, 2011; Kaneko-Suzuki *et al.*, 2018). Based on *FT1* transcripts being higher in photoperiod-insensitive lines and spikelet identity genes being expressed at lower levels in *ft-b1* mutants, we hypothesised meristem identity genes would be more highly expressed in *Ppd-D1a* NILs, relative to wild-type (Beales *et al.*, 2007; Shaw *et al.*, 2013; Boden *et al.*, 2015). Surprisingly, while photoperiod-insensitive lines accelerated meristem identity gene expression to occur earlier in the season, the overall amplitude of transcripts was identical to wild-type. In *ppd-1* lines, induction of meristem identity genes was delayed, and transcript levels were lower than those detected in wild-type, consistent with the reduced activity of *FT1* in these lines. The step-wise increase in *FT1* expression in leaves aligned strongly with the seasonal up-regulation of meristem identity genes and progression of IM development, with the first induction of *FT1* facilitating the vegetative to double-ridge transition and the second rise promoting advancement to later stages (**Fig. 7**). The accelerated peak in meristem identity gene expression in photoperiod-insensitive lines coincided with arrival of IMs at the terminal spikelet stage earlier in the season. These data indicate that the regulation of inflorescence development and spikelet number by *Ppd-1* is not determined by the absolute level of meristem identity gene expression, but by the timing at which the peak occurs. The advanced induction of *FT1* in the photoperiod-insensitive line potentially explains its ability to advance to terminal spikelet when daylengths were maintained at 10 h, relative to wild-type and *ppd-1* that stalled at the lemma primordium stage. In addition to regulating spikelet number, photoperiod-insensitive alleles also reduce floret fertility; the increased expression of *GNI1* at later stages in *Ppd-D1a* NILs, relative to wild-type, may explain the decrease in fertile florets (Prieto *et al.*, 2018; Sakuma *et al.*, 2019). Taken together with the seasonal analysis of *FT1* expression, these results indicate that inflorescence development is intimately connected with the activity of floral signals generated in leaves, which dynamically respond to increasing daylengths.

Our analysis identified *FT2* as a key regulator of spikelet development in hexaploid wheat. Based on the *ft-b2* mutants producing more spikelets, and the significant increase in *FT2* expression between the lemma primordium and terminal spikelet stages, we propose *FT2* helps determine spikelet number by promoting transition of the IM to the terminal spikelet stage. This conclusion is consistent with analysis performed in tetraploid wheat, which showed *FT2* is expressed strongly in the developing IM and that loss-of-function *ft2* alleles delay flowering and increase spikelet number (Shaw *et al.*, 2019). A role for *FT2* in spikelet termination is supported by transcript analysis in *ppd-1* and photoperiod-insensitive *Ppd-D1a* NILs, in which *FT2* expression was inversely proportional to spikelet number and the rate of inflorescence development. The early rise of *FT2* expression in photoperiod-insensitive *Ppd-D1a* NILs may explain why IMs from this line could progress to the terminal spikelet stage when plants were maintained at 10 h photoperiods, relative to wild-type, in which *FT2* expression was significantly lower than those grown under natural photoperiods. Interestingly, the rise in *FT2* transcripts between the lemma primordium and terminal spikelet stages and its genetic association with spikelet number contrasts the profile of genes such as *AP1, SEP1, VRN1* and *WFZP*, for which expression increases earlier in IM development and is associated with supernumerary spikelet formation (Boden *et al.*, 2015; Dobrovolskaya *et al.*, 2015). These results indicate that genes induced between the lemma primordium and terminal spikelet stages contribute to determination of spikelet number, while those expressed at earlier stages influence inflorescence architecture. Together with studies that have investigated the inflorescence transcriptome of wheat, these results allude to a strategy for identifying genes that regulate spikelet number (Wang *et al.*, 2017; Li *et al.*, 2018). For example, expression of *TERMINAL FLOWER 1* (*TFL1*) and *WHEAT ORTHOLOGUE OF APO1* correlates with spikelet number, with increased transcripts during the glume and floret primordium stages being associated with extra spikelets (Wang *et al.*, 2017; Kuzay *et al.*, 2019).

In summary, our work provides new insights into the regulation of molecular processes controlling flowering and inflorescence development in the field. We show that the early stages of inflorescence development are coordinated by step-wise inductions of *FT1* expression, which are overridden by photoperiod-insensitive *Ppd-D1a* alleles. The results highlight the importance of complementing laboratory-based analysis with experiments performed in the field – a concept echoed by work performed in oilseed rape, Arabidopsis and rice that have used field-grown plants to uncover new information about the molecular events controlling seasonal regulation of flowering (Duncan *et al.*, 2015; Gomez-Ariza *et al.*, 2015; Hepworth *et al.*, 2018; Song *et al.*, 2018; O’Neill *et al.*, 2019). Our work provides an important foundation for understanding the mechanisms controlling yield-related traits of wheat, which is vital given our increasing need to improve global food security by generating superior yielding cultivars (Fischer *et al.*, 2014).

## ACKNOWLEDGEMENTS

We acknowledge the BBSRC Norwich Research Park Doctoral Training Partnership for A.G. (BB/M011216/1), the BBSRC Designing Future Wheat programme (BB/P016855/1) and the Royal Society (UF150081) for funding the research. We thank Simon Griffiths, Richard Morris and Laura Dixon for helpful comments during the project, and the horticultural and field services team at JIC for assistance with plant husbandry.

## AUTHOR CONTRIBUTIONS

A.G. and S.A.B. contributed to conceptualization, formal analysis, methodology, investigation, validation, visualization and writing (original draft preparation and review/editing). S.A.B. also contributed to supervision, project administration and funding acquisition.

