## Supplementary information for "Step-wise increases in *FT1* expression regulate seasonal progression of flowering in wheat (*Triticum aestivum* L.)"

***New Phytologist*** **Supporting Information**

Article title: **Step-wise increases in *FT1* expression regulate seasonal progression of flowering in wheat (*Triticum aestivum* L.)**
Authors: Adam Gauley and Scott A. Boden
Article acceptance date: *tbd*

### Supplementary Figures

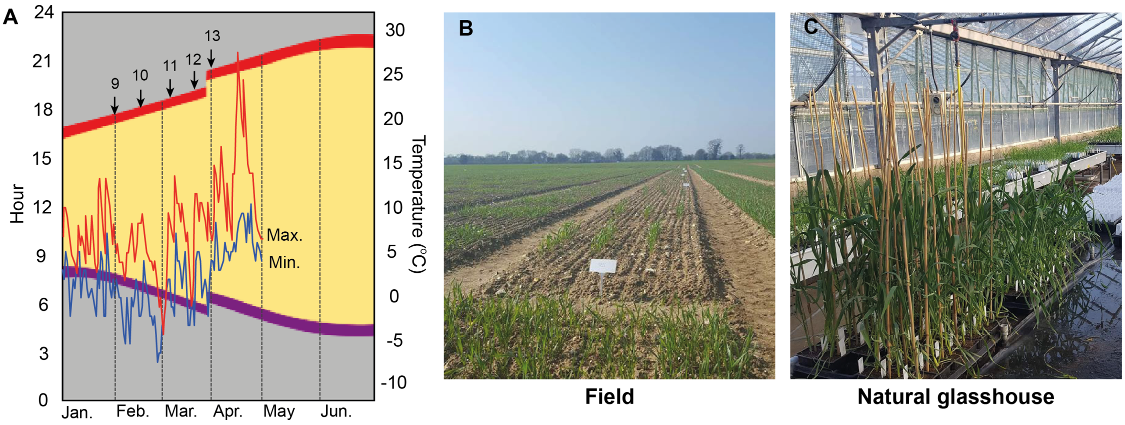


**Figure S1: Seasonal events of photoperiod experiment in Norwich, UK. (A)** Schematic of seasonal changes in daylength as winter transitions into spring and summer, with periods of light (yellow) and night (grey) that occur in Norwich (52°62'25.7"N, 1°21'83.2"E), highlighting collections points for each hourly increase in daylength. Maximum (red line) and minimum (blue line) temperatures are shown for the duration of the experiment performed in 2019. Images of the experimental site for the field- **(B)** and glasshouse-based **(C)** experiments.

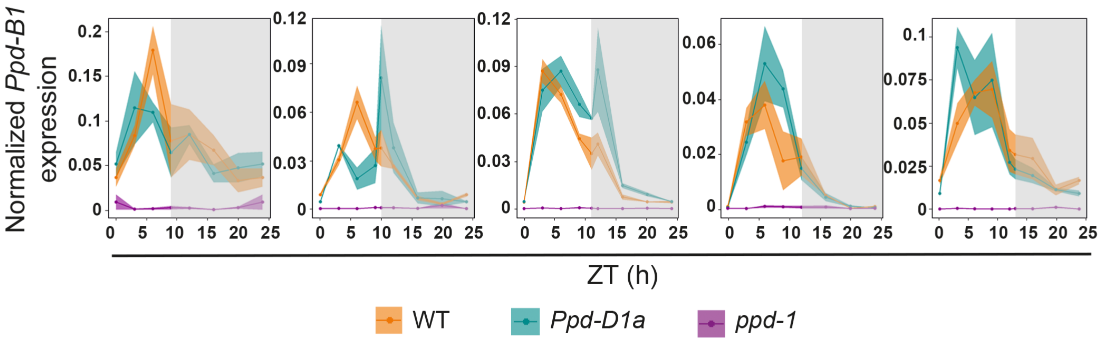


**Figure S2: Seasonal regulation of *Ppd-B1* in the field.** Diurnal expression profiles of *Ppd-B1* in wild-type (yellow), *Ppd-D1a* photoperiod-insensitive (green) and *ppd-1* (magenta) NILs under field conditions at different photoperiods. The grey shading represents night-time hours. All expression profiles are shown over a 24-hour period at hourly incremental increases in daylength. Each point is the normalized mean transcript levels ± SEM of three biological replicates.

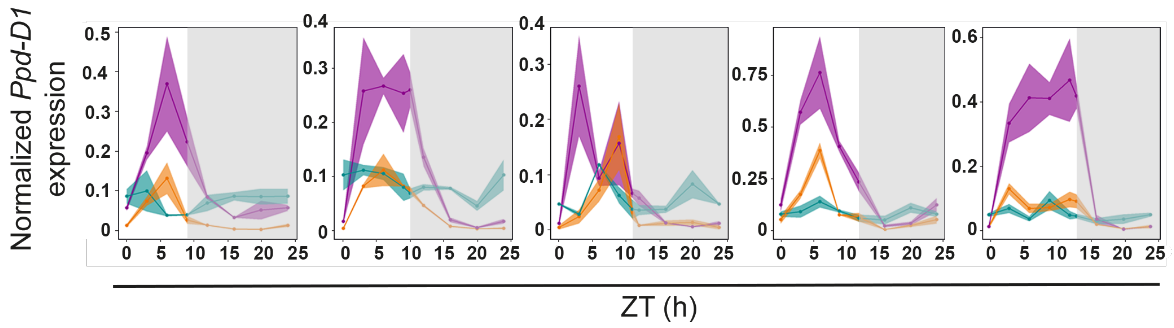


**Figure S3: Seasonal regulation of *Ppd-D1* in the glasshouse.** Diurnal expression profiles of *Ppd-D1* in wild-type (orange), *Ppd-D1a* photoperiod-insensitive (cyan) and *ppd-1* (magenta) NILs under field conditions at different photoperiods. The grey shading represents night-time hours. All expression profiles are shown over a 24-hour period at hourly incremental increases in daylength. Each point is the normalized mean transcript levels ± SEM of three biological replicates.

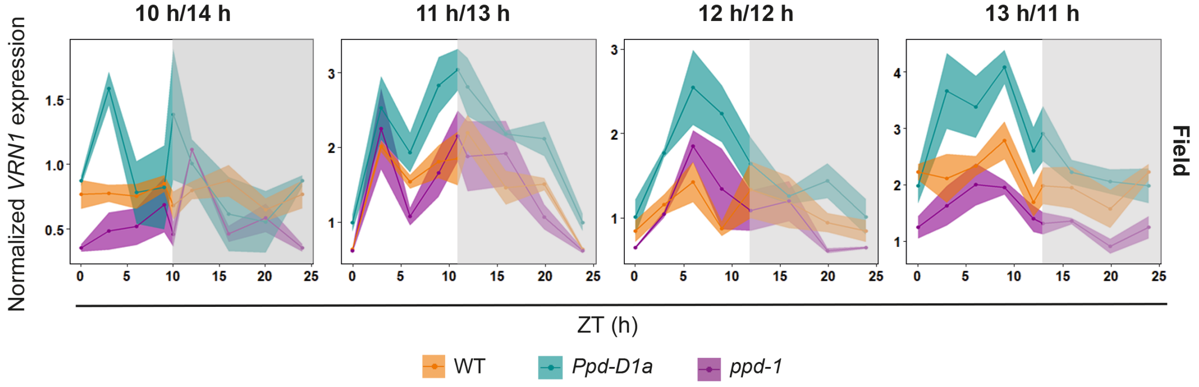


**Figure S4: Seasonal regulation of *VRN1* expression in the field.** Diurnal expression profile of *VRN1* in wild-type (orange), *Ppd-D1a* photoperiod-insensitive (cyan) and *ppd-1* (magenta) NILs under field conditions. The grey shading represents night-time hours. Each point is the normalized mean transcript levels ± SEM of three biological replicates.

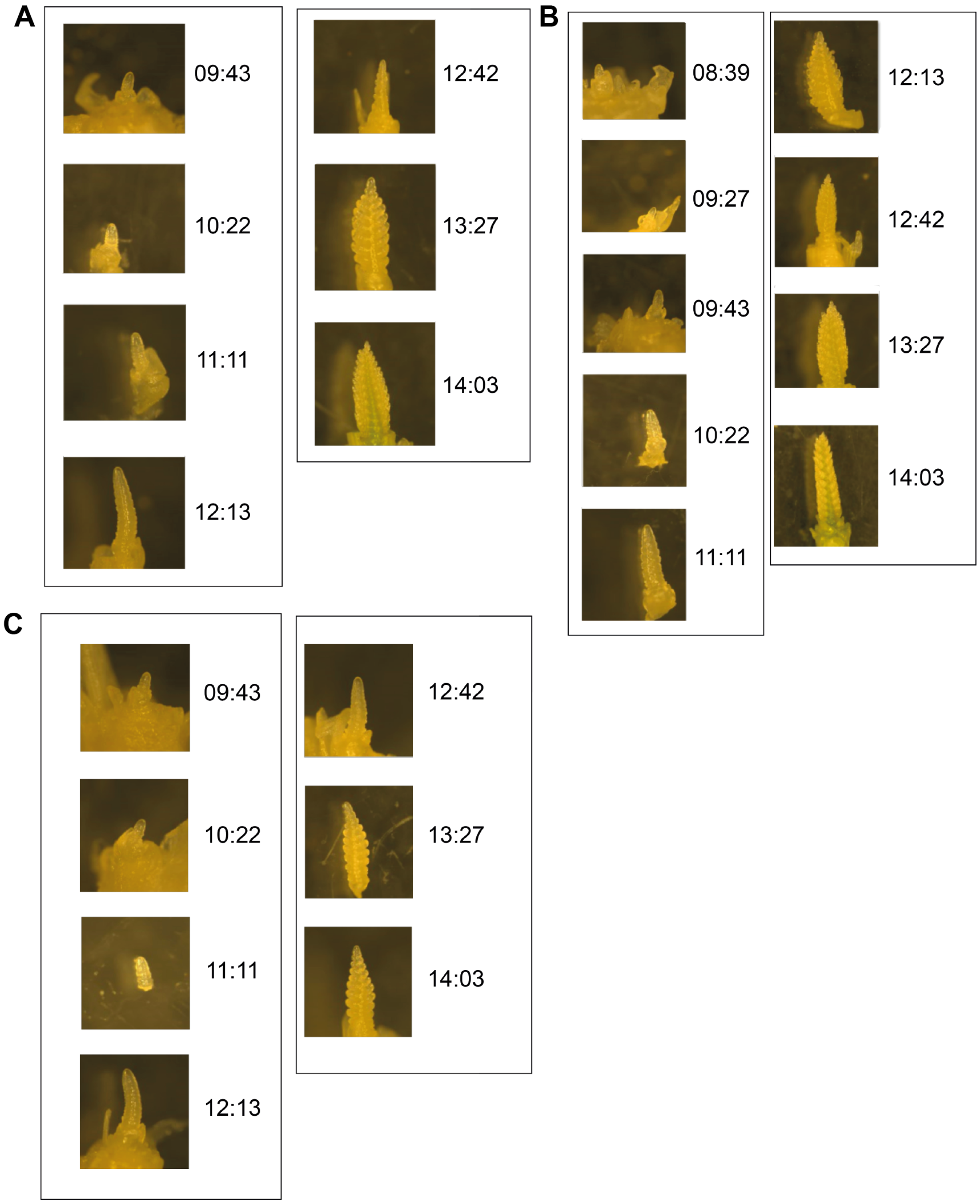


**Figure S5: Seasonal progression of inflorescence meristem development.** Representative photographs of inflorescence meristems from (**A**) wild-type, (**B**) *Ppd-D1a* photoperiod-insensitive and (**C**) *ppd-1* NILs of field-grown plants. Corresponding daylengths (h:min) are shown. Please note that scale bars are not shown as images were taken at different magnitudes according to stage – these images are shown to detail progression of developmental stage in comparison to the other two genotypes.

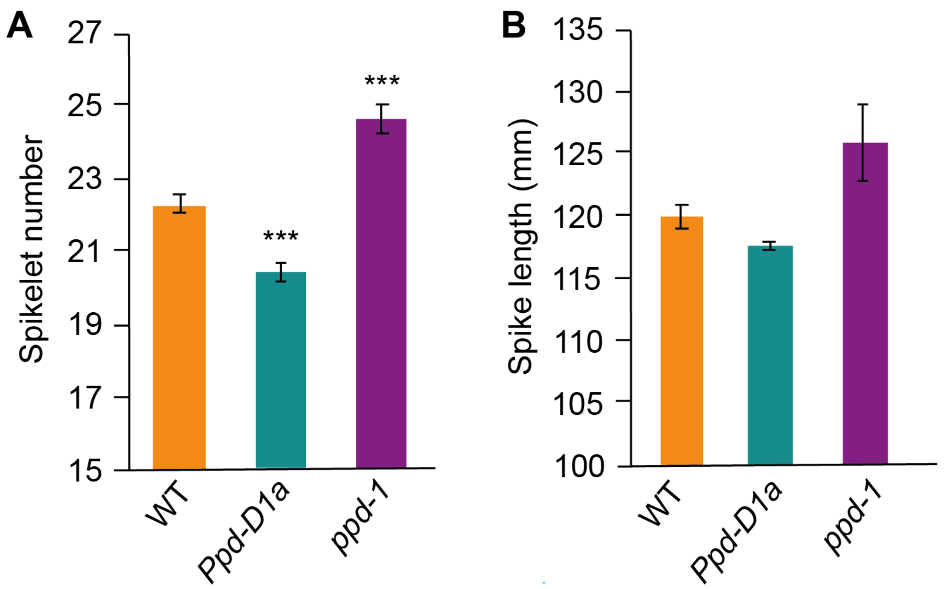


**Figure S6: Inflorescence architecture phenotype of *Ppd-1* lines. (A)** Inflorescence length phenotypes for wild-type, *Ppd-D1a* and *ppd-1* NILs grown under field conditions. Data are the mean ± SEM of 10 biological replicates

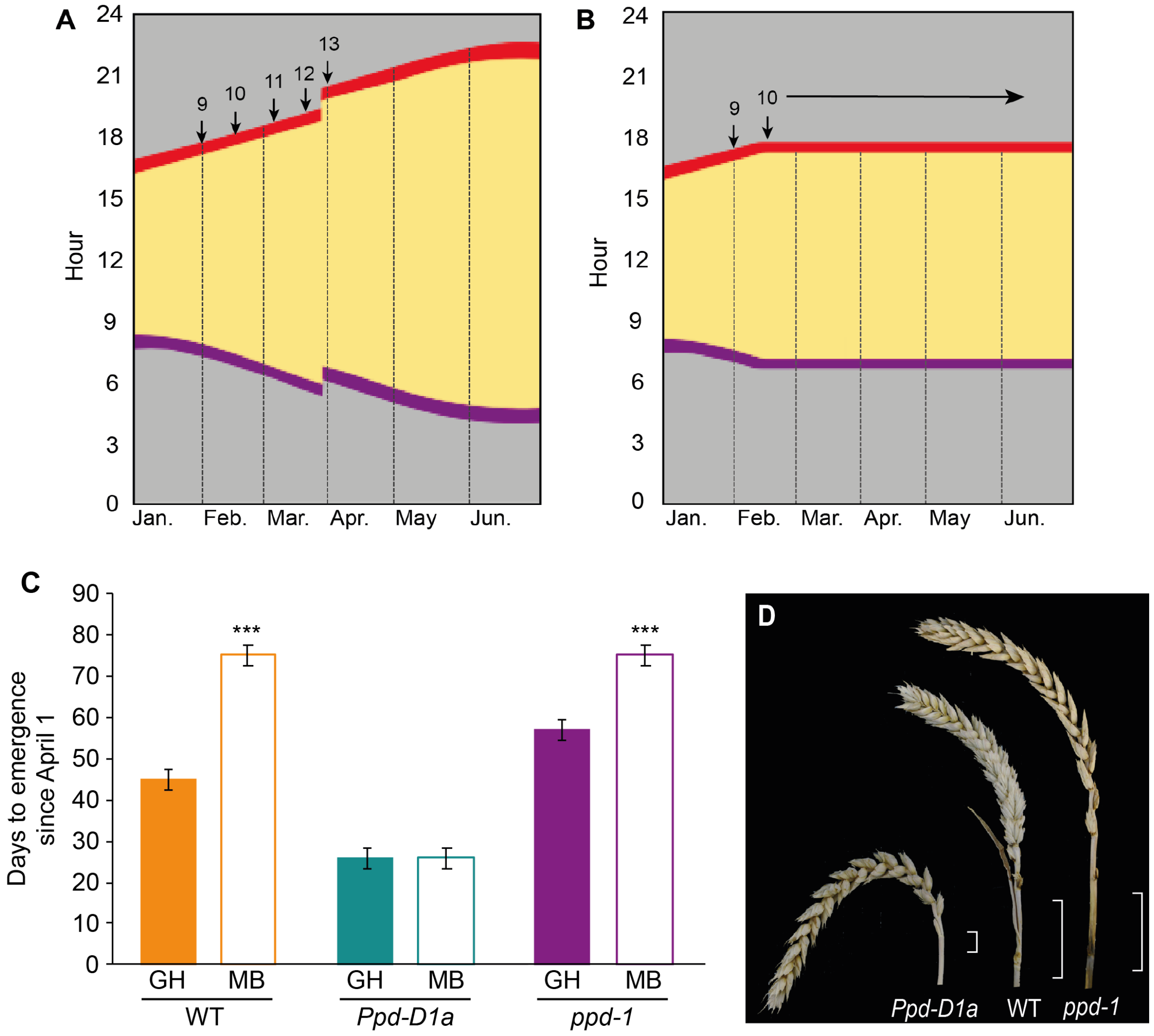


**Figure S7: Experimental design and phenotypes of plants from the photoperiod shift experiment.** Schematic diagrams of seasonal changes in daylength as for plants that were **(A)** maintained under natural photoperiods, relative to **(B)** plants that were grown under natural photoperiods until daylengths were 10 h, and then maintained under 10 h using a glasshouse equipped with a moving bench. (**C**) Flowering time measurements of wild-type (WT, orange), *Ppd-D1a* NIL (green) and *ppd-1* null NILs grown under natural photoperiod glasshouses (GH, solid) or the moving bench conditions (MB, white). (**D**) Images of inflorescences from plants of all three genotypes grown under MB conditions, showing elongated internodes (white brackets) Data are mean ± SEM of five biological replicates.

### Supplementary Methods

The following methods are written as supplementary information due to word limits in the main text.

#### *Inflorescence meristem phenotyping*

Stages of inflorescence development were determined as per (Kirby & Appleyard, 1987). Plants from the glasshouse and field were isolated and imaged using a Leica MZ16 Stereo binocular dissecting microscope and a Lecia DFC420 colour camera for imaging. Each data point included 4 - 5 biological replicates.

#### *Heading date measurements*

Heading dates for all field- and glasshouse-grown plants were measured as the calendar date of flowering. For field-grown plants, heading date was determined per plot on the day that 50% of the ear had emerged from 50% of plants, with data the average ± SEM of 5 replicates. For glasshouse- and moving bench-grown plants, heading date was recorded for individual plants. Data are average ± SEM of 20 and 7 replicates, respectively. Heading-date was defined as the day when half of the inflorescence had emerged from the sheath on the main tiller.

#### *DNA extractions and sequence analysis*

Genomic DNA extractions were carried out as described previously (Dixon *et al.*, 2018). All gene sequences were obtained by BLAST search (Ensembl Plants). Primers used for *ft-b2* mutant alleles are listed in Table S1. DNA fragments were amplified using Phusion DNA polymerase (New England Biolabs). Amplicon sequencing was carried out with Mix2Seq Kit (Eurofins).

#### *Kompetitive Allele-Specific PCR analysis*

Kompetitive allele-specific PCR (KASP) analysis was used to analyse TILLING mutant lines, prior to verification of mutations using sequence analysis. Oligonucleotides were designed using Polymarker (Ramirez-Gonzalez *et al.*, 2015) and they contained either the standard FAM or HEX compatible tails (Table S2). The assay was performed as described previously (Dixon *et al.*, 2018).

#### *Statistical analysis*

Statistical differences between treatments and sample points were tested by two-tailed Students *t* test. Data in figures are mean ± standard error of the mean (SEM). All outcomes of the statistical analysis relating to gene expression are provided in Supplementary Dataset 1.

**Table S1: Sequences of oligonucleotides used in qRT-PCR assays.**

| Gene | Direction | Primer | Source |
|---|---|---|---|
| *SEP1-6* | Forward | CCTCTACCAGTTCTCCTCCTCC | Boden et al. 2015 |
| | Reverse | CATATACTCCAGATAGTTGTT | previously *AGLG1* |
| *VRN1* | Forward | GGAAACTGAAGGCGAAGGTTGA | Oliver et al. 2013 |
| | Reverse | TGGTTCTTCCTGGATCTGATATG | |
| *AP1-3* | Forward | TCTATGAGTACGCCACCGACT | Boden et al. 2015; |
| | Reverse | CACCAATTTCCCTCACTTTCA | previously *AGL29* |
| *AP1-2* | Forward | AGCTCACCGTCACCTACACC | Boden et al. 2015 |
| | Reverse | TTGTTTGCTTGTGCTGGAGA | previously *AGL10* |
| *AGL6* | Forward | CCAGACAGCGAAAGACACAA | This paper |
| | Reverse | CTTGTGCTTGAGTTGCCTGT | |
| *FT1* | Forward | GTCGTTCGGGCAGGAG | Shaw et al. 2012 |
| | Reverse | TGGAAGAGTACGAGCACGA | |
| *Ppd-D1* | Forward | AAGACAAGGCTGATGAAATGAG | Shaw et al. 2012 |
| | Reverse | GAAGGATTGACCACATTGGA | |
| *Ppd-B1* | Forward | AAGACAAGGTTGATGACGTGA | Shaw et al. 2012 |
| | Reverse | GAGGGATTGATCACGTTGG | |
| *SOC1* | Forward | CAGCAAGTCAAAGCTGATGC | Boden et al. 2015 |
| | Reverse | AACGCGGAGACTCTTCTCAA | |
| *GNI1* | Forward | AGCTTATGGAGGAGGAGTTCG | This paper |
| | Reverse | CTCCCAGCCTCTCCTTCAG | |
| *FT2* | Forward | GCTGGTCACCGACATCCCGG | This paper |
| | Reverse | CCTCGTAGCACACCACCTCG | |

**Table S2: Sequences of oligonucleotides used for genotype analysis of *FT-B2* mutants**

| Gene | Direction | Mutant line | Primer |
|---|---|---|---|
| *FT-B2* | KASP | Cadenza122 | gaaggtgaccaagttcatgctGGACGGCCTCAGCTCGCAGC |
| *FT-B2* | KASP | Cadenza122 | gaaggtcggagtcaacggattGGACGGCCTCAGCTCGCAGT |
| *FT-B2* | KASP | Cadenza122 | CGGCCGCCTTCATTAACATA |
| *FT-B2* | Forward | Cadenza1655 | GAGGTGGTGTGCTACGAGG |
| *FT-B2* | Reverse | Cadenza1655 | CCCATCTGATTCCCGTACGA |
